## supplementary text and figs for "A chemically-induced attenuated strain of *Candida albicans* generates robust protective immune responses and prevents systemic candidiasis development"

**<u>Supplementary Informations</u>**

**Supplementary Figure S1. Identification of a fungistatic concentration of EDTA. (A)** Various concentrations of EDTA ranging from 0.9 µM to 500 µM was added to the *C. albicans* culture with a total volume of 150 µL in a 96-well plate and allowed to grow overnight at 30 °C under static condition. YPD media alone and without EDTA treatment culture as controls were also taken. After 24 hrs, the image of the 96-well polystyrene plate was taken. **(B)** After 24 hr, the absorbance was measured at $OD_{600\ nm}$ using a Perkin-Elmer plate reader and plotted using GraphPad Prism software version 8.0. *p* value as indicated ****< 0.0001 was determined by one-way ANOVA. **(C)** *C. albicans* cells treated without and with 62.5 µM, 125 µM, and 250 µM concentration of EDTA for 24 hrs and indicated dilutions were spotted on YPD-agar plate. The plate was incubated at 30 °C for 24 hrs and imaged using a Chemidoc imaging system.

**Supplementary Figure 2: Effect of 250 µM EDTA on cell viability.** *C. albicans* pre-culture was diluted to an $OD_{600nm}$=0.5 and allowed to grow in the absence and presence of 250 µM EDTA at 30 °C and 200 rpm up to 12 hrs. (**A**) At mentioned time points (0 -12 hrs), cells were harvested, stained with SYTOX Green and analyzed by flow cytometry (blue laser; excitation at 488nm). The data acquisition was performed using the BD LSR Fortessa Flow cytometer and analysis was carried out by FlowJo version 8.1 software. Analysed data exported in JPEG format. Acquisition profile of unstained cells was also shown. (**B**) The percentage of dead cells population from two technical replicates was plotted using GraphPad Prism software version 8.0. No statistical significance difference was found between treated and untreated data. (**C**) Similarly, the harvested cells were stained with PI and images were captured using a fluorescence microscope (EVOS imaging system; Thermo Fisher Scientific) at 40X magnification and the red dots indicated the dead cells. (**D**) The population of dead cells was determined from 15 microscopy frames for each time interval and plotted using GraphPad Prism. *p* value that was non-significant as determined by two-way ANOVA.

**Supplementary Figure 3: Effect of EDTA on *C. albicans* morphology.** *C. albicans* pre-culture was diluted to an $OD_{600nm}$=0.5 and allowed to grow in the absence and presence of 250 µM EDTA at 30 °C and 200 rpm up to 12 hrs. Similar experiment was also carried with 10% serum and morphological transition was induced at 37 °C. (**A**) At various time intervals**,** cell morphology was observed using a 40X Leica DM500 microscope (scale bar of 2 µm). (B) The percentages of singlet, doublet, pseudohyphae, and hyphae cells were quantified without (i) and with serum (ii).

(C) The length of serum induced germ tubes (n=60) in µm was measured using ImageJ software. *p* value as indicated **<0.01 or non-significance as determined by two-way ANOVA.

**Supplementary Figure S4. Transcriptomic analyses of CAET and validation. (A)** Principal Component Analyses (PCA) was performed for proper grouping of three biological replicates represented as MT523, MT524, and MT525 as one cluster for *Ca* and MT526, MT527, and MT528 for CAET for another. The plot scores indicate the principal component 1 vs. principal component 2 signifying the largest variable vs. second largest variable. **(B)** Gene expression was performed using qRT-PCR and semi-quantitative assay for selected upregulated genes obtained from the heat map i.e., **(i)** metal ion associated gene *ZRT1*, **(ii)** cell wall associated gene *CHT4* (chitin-specific gene), and **(iii)** host-pathogen interaction-related gene *PGA13*. Similarly, the downregulated genes selected are **(iv)** aconitase *ACO2*, **(v)** virulence-associated gene *PLB1*, and **(vi)** thiamine synthesis-associated gene *THI13* expression were analysed. *GAPDH* considered as control. *p* values *< 0.05 and ****< 0.0001 were as determined by unpaired t test. **(C)** STRING analysis showing only the clusters related to 40S (i), 60S subunits (ii) and one-carbon metabolism (iii). An interaction score of 0.7 was considered for all the networks. (**D**) Gating strategy of unstained RAW cells alone was shown.

**Supplementary Figure S5. Systemic candidiasis development in *C. albicans* challenged mice**. Mice (n=6/category) were injected intravenously with 5 x $10^5$ *C. albicans* cells (untreated *Ca*: cyan blue and CAET: purple) and saline as control (grey) and their survivability was monitored for 30 days. In a similar set of experiment, mice (n=6/category) were first immunized with CAET (represented in brown) or sham vaccinated with saline (represented in orange), and after 30 days, they were further re-challenged with *C. albicans*. Their survivability was monitored for another 30 days and a survival curve was plotted. 1° and 2° suggest primary and secondary challenges, respectively. Mice suffered due to severe infections were euthanized and fungal load in vital organs were determined by CFU analyses. Statistical significance of the survival curve was determined using Mantel-Cox test. *p* values were as mentioned and others were non-significant. Y-axis was represented in Log rank scale. Two repeat studies were shown.

**Supplementary Figure S6. (A)** PAS-stained kidney sections of different mice groups (saline, *Ca*, CAET and 1°CAET 2°*Ca*) from the experiment in Fig. 5D showing fungal burden at different times of sacrifice post inoculation. (**B**) Lymphocytes, RBC, MCHC, MCH, and MCV (fl) present in total blood of various mice groups sacrificed on various days from the experiment in Fig. 5D was estimated. *p* values *< 0.05, **<0.01, ***<0.001, and ****< 0.0001 as determined by two-way ANOVA.

**Table S1**: List of top 100 upregulated genes in EDTA treated *C. albicans* cell (CAET) with detailed information

**Table S2**: **(A)** GO annotation analysis for 411 upregulated genes and **(B)** 388 downregulated genes.

**Table S3**: **(A)** STRING cluster analysis for 411 upregulated genes, **(B)** 388 downregulated genes, and **(C)** 74 downregulated genes out of 388 DEGs specific to 40S and 60S ribosomal subunits.

**Table S4**: STRING cluster analysis for 33 DEGs specific to pathogenesis.

**Table S5** List of top 100 downregulated genes in EDTA treated *C. albicans* cell (CAET) with detailed information

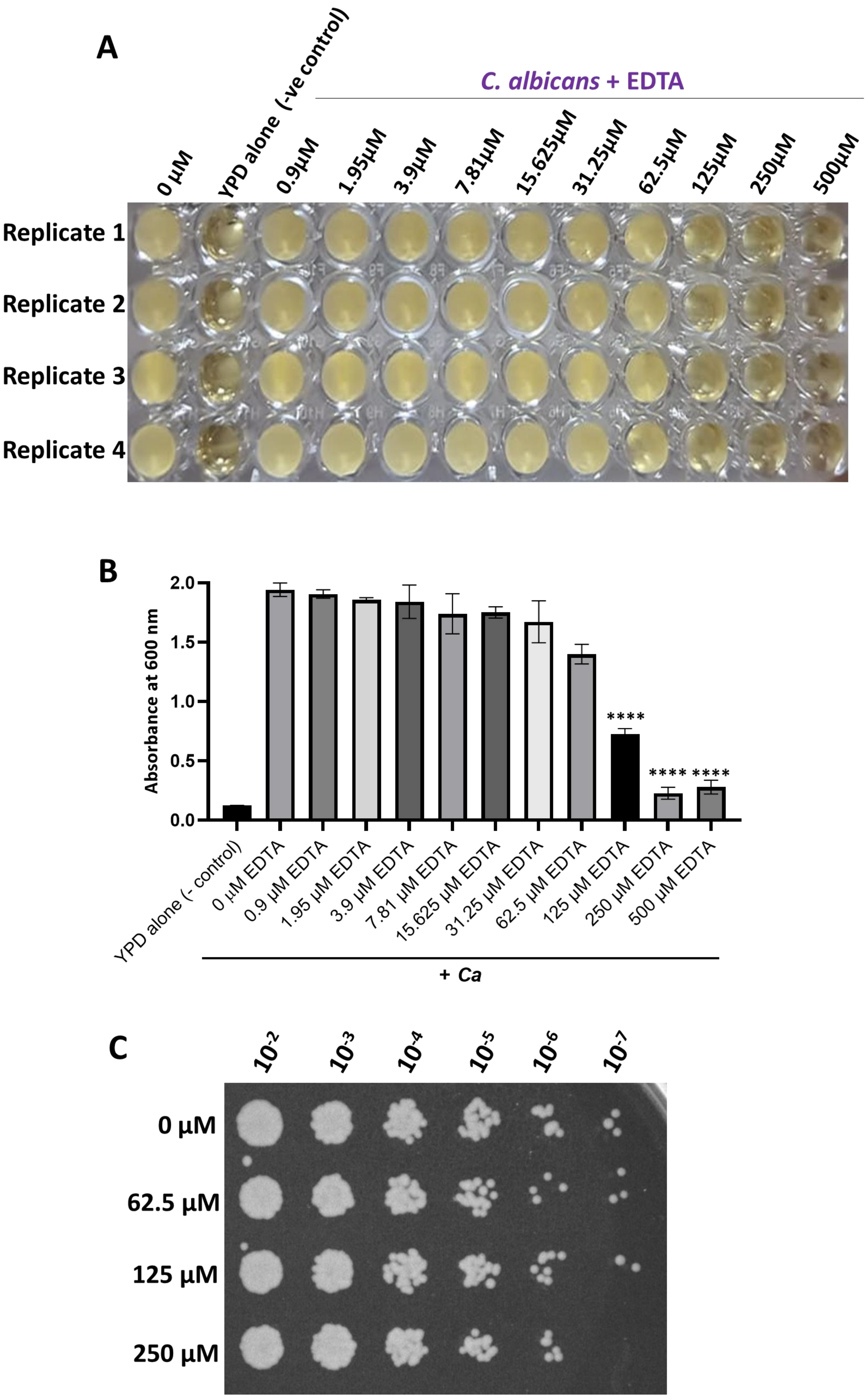
A
C. albicans + EDTA
0 µM
YPD alone (-ve control)
0.9µM
1.95µM
3.9µM
7.81µM
15.625µM
31.25µM
62.5µM
125µM
250µM
500µM
Replicate 1
Replicate 2
Replicate 3
Replicate 4
B
Absorbance at 600 nm
2.0
1.5
1.0
0.5
0.0
****
**** ****
YPD alone (- control)
0 µM EDTA
0.9 µM EDTA
1.95 µM EDTA
3.9 µM EDTA
7.81 µM EDTA
15.625 µM EDTA
31.25 µM EDTA
62.5 µM EDTA
125 µM EDTA
250 µM EDTA
500 µM EDTA
+ Ca
C
10-2
10-3
10-4
10-5
10-6
10-7
0 µM
62.5 µM
125 µM
250 µM

Figure S1

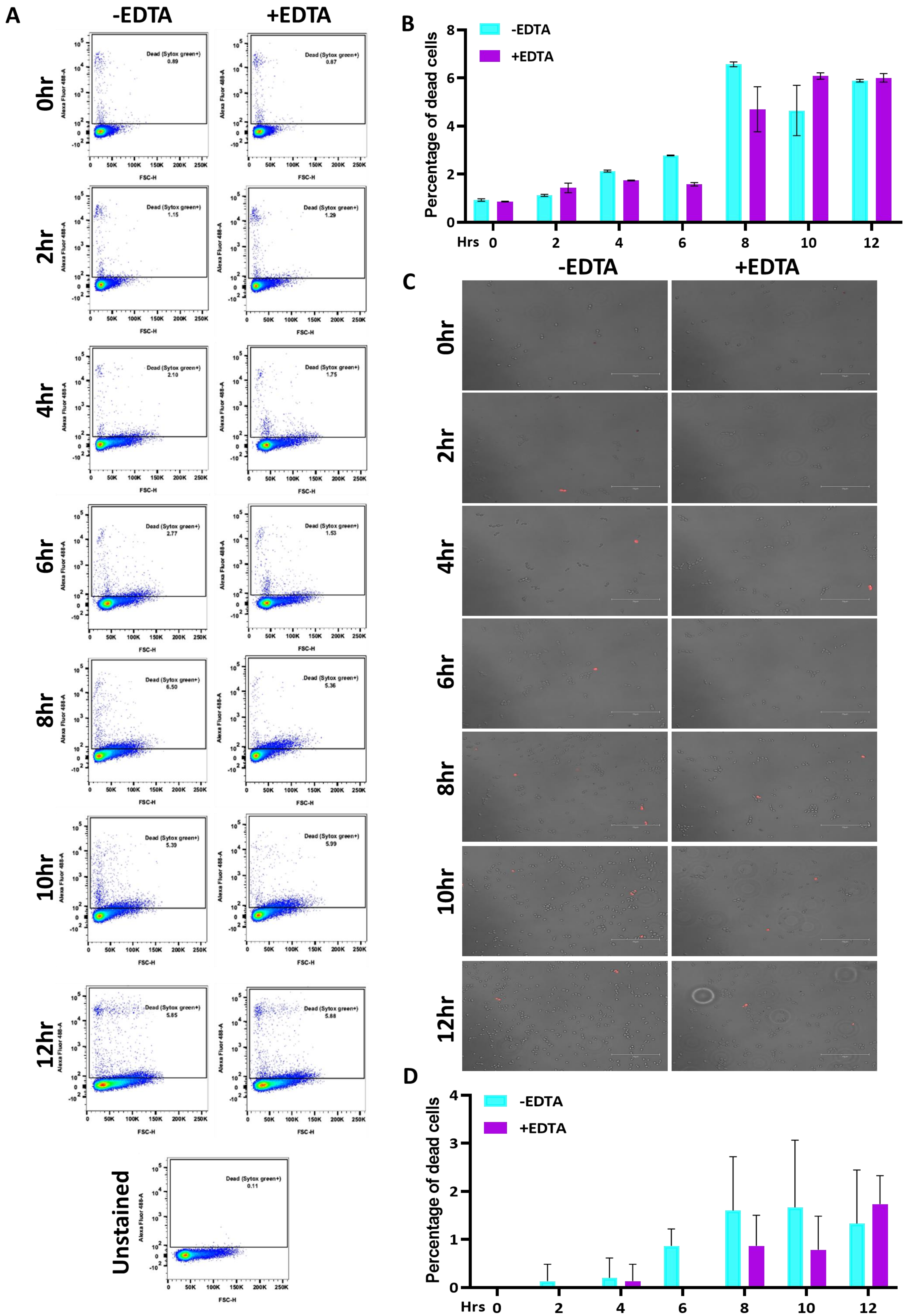

A
-EDTA
+EDTA
0hr
2hr
4hr
6hr
8hr
10hr
12hr
Unstained
Dead (Sytox green+)
Alexa Fluor 488-A
FSC-H
B
Percentage of dead cells
-EDTA
+EDTA
Hrs
C
D


Figure S2

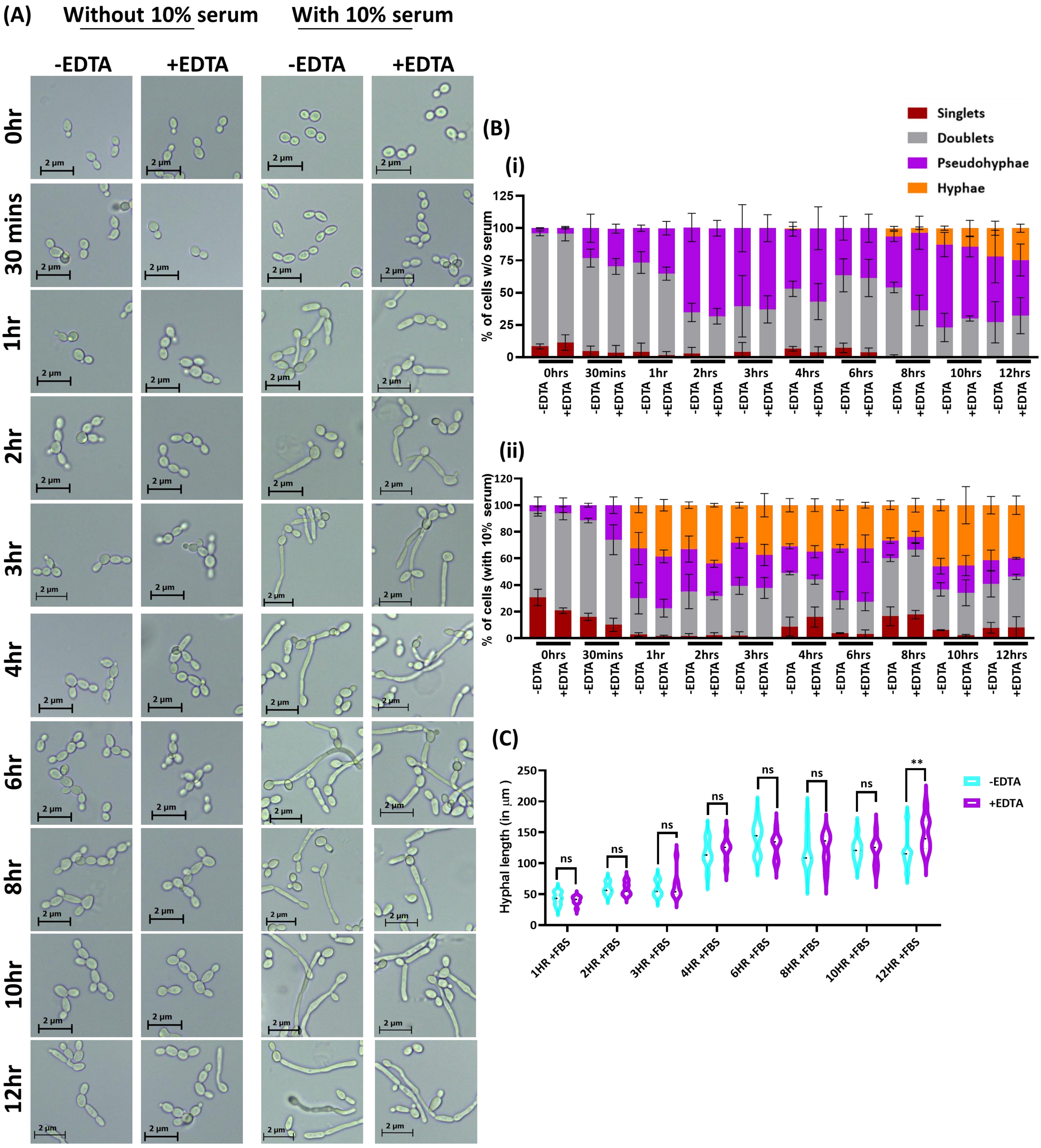
(A)
Without 10% serum
With 10% serum
-EDTA
+EDTA
-EDTA
+EDTA
0hr
30 mins
1hr
2hr
3hr
4hr
6hr
8hr
10hr
12hr
2 µm
(B)
(i)
Singlets
Doublets
Pseudohyphae
Hyphae
% of cells w/o serum
0hrs
30mins
1hr
2hrs
3hrs
4hrs
6hrs
8hrs
10hrs
12hrs
-EDTA
+EDTA
(ii)
% of cells (with 10% serum)
(C)
Hyphal length (in µm )
ns
**
-EDTA
+EDTA
1HR +FBS
2HR +FBS
3HR +FBS
4HR +FBS
6HR +FBS
8HR +FBS
10HR +FBS
12HR +FBS

Figure S3

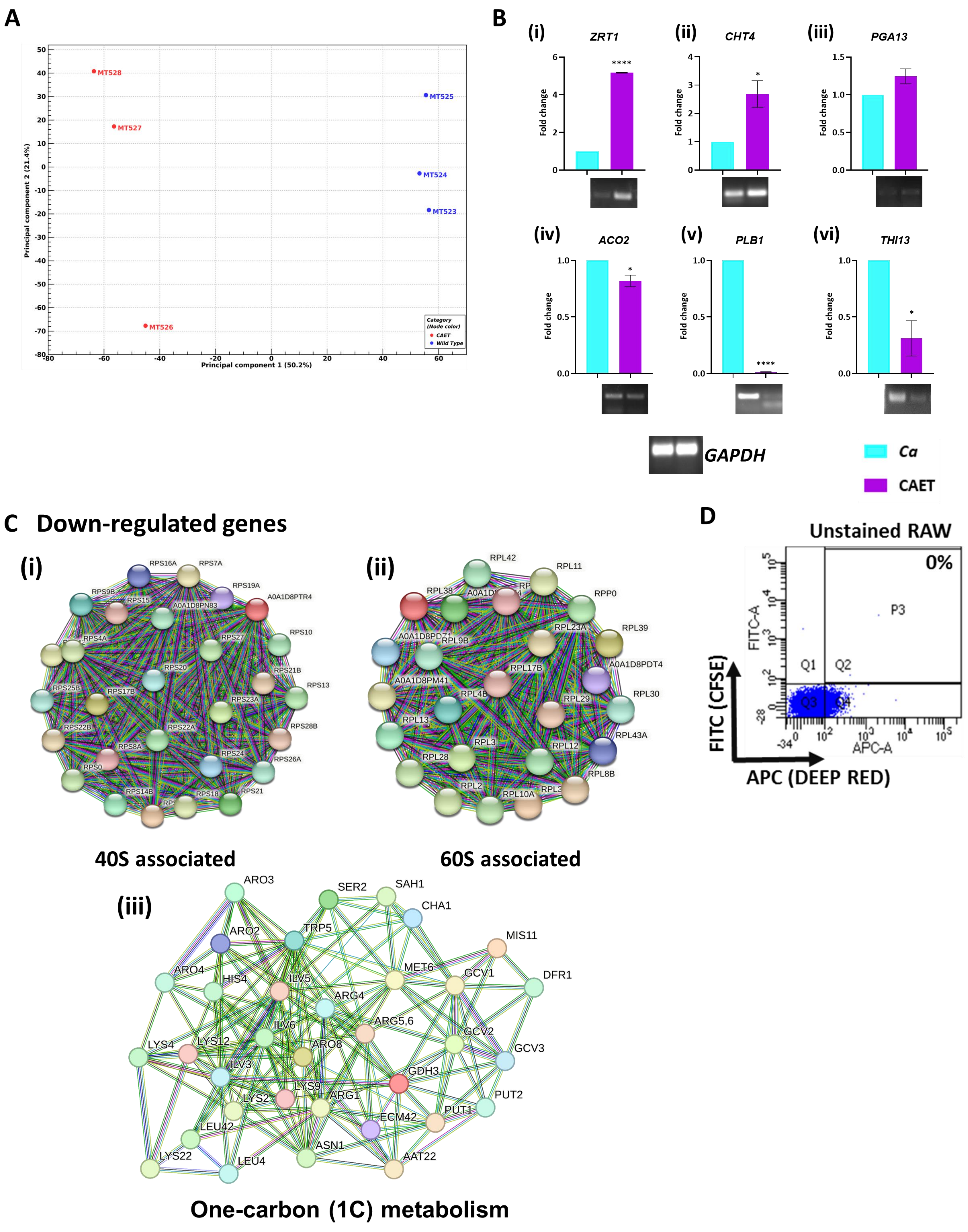
A
Principal component 1 (50.2%)
Principal component 2 (21.4%)
MT528
MT527
MT526
MT525
MT524
MT523
Category (Node color)
CAET
Wild Type
B
(i) ZRT1
(ii) CHT4
(iii) PGA13
(iv) ACO2
(v) PLB1
(vi) THI13
Fold change
GAPDH
Ca
CAET
C Down-regulated genes
(i)
(ii)
40S associated
60S associated
(iii)
One-carbon (1C) metabolism
D
Unstained RAW
0%
FITC (CFSE)
APC (DEEP RED)

Figure S4

### Repeat 1

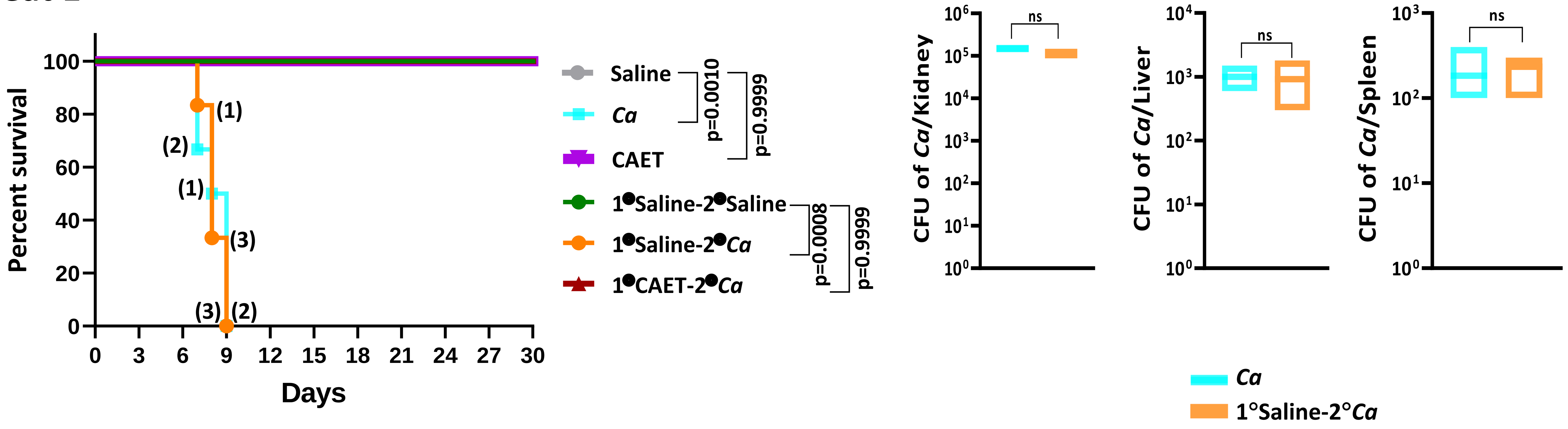


### Repeat 2

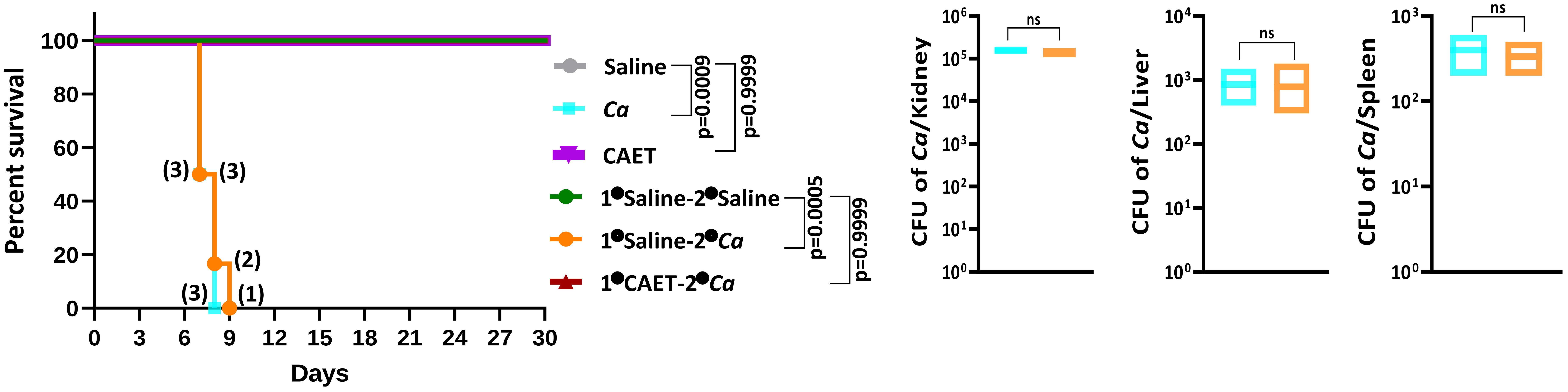


**Figure S5**

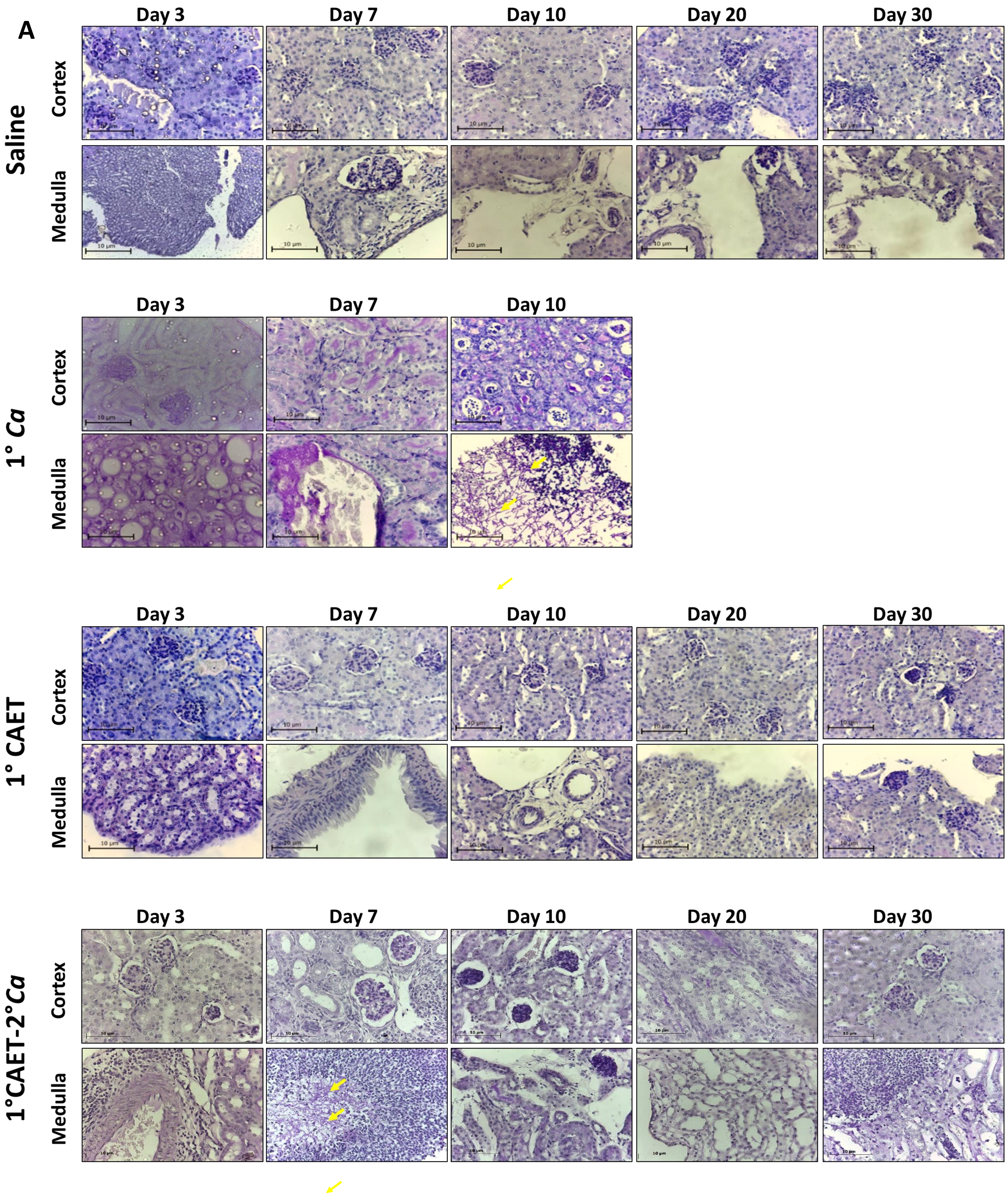
A
Saline
Cortex
Medulla
Day 3
Day 7
Day 10
Day 20
Day 30
1° Ca
1° CAET
1°CAET-2°Ca
10 μm

Figure S6

B

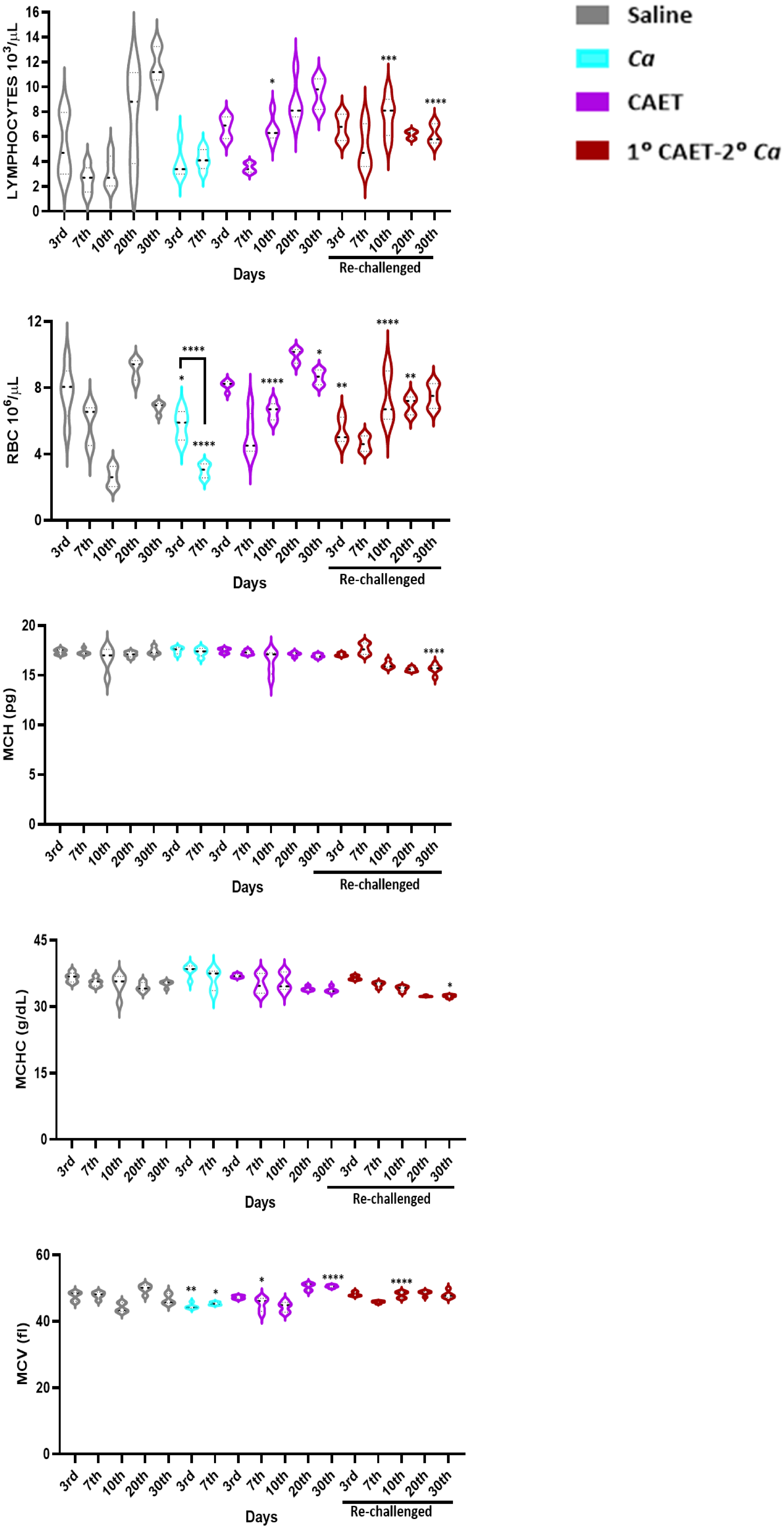

Saline
Ca
CAET
1° CAET-2° Ca
LYMPHOCYTES 10³/µL
RBC 10⁶/µL
MCH (pg)
MCHC (g/dL)
MCV (fl)
3rd
7th
10th
20th
30th
Days
Re-challenged
*
**
***
****


Figure S6
